# Hematopoietic progenitors polarize in contact with bone marrow stromal cells by engaging CXCR4 receptors

**DOI:** 10.1101/2020.05.11.089292

**Authors:** Thomas Bessy, Benoit Souquet, Benoit Vianay, Alexandre Schaeffer, Thierry Jaffredo, Jerome Larghero, Laurent Blanchoin, Stephane Brunet, Lionel Faivre, Manuel Théry

**Affiliations:** Cytomorpho Lab, HIPI, U976, INSERM / CEA / AP-HP / Université de Paris, Institut de Recherche Saint Louis, Paris, France; Cytomorpho Lab, LPCV, UMR5168, CEA / INRA / CNRS / Univ. Grenoble-Alpes, Interdisciplinary Research Institute of Grenoble, Grenoble, France; Alveole, 68 Boulevard de Port-Royal, 75005, Paris, France; Laboratoire de Biologie du Développement, CNRS UMR 7622, Inserm U1156, Sorbonne Université, Institut de Biologie Paris-Seine, Paris, France; AP-HP, Hôpital Saint-Louis, Unité de Thérapie Cellulaire; Inserm U976; Université de Paris

## Abstract

Hematopoietic stem and progenitor cells (HSPCs) are located in the bone marrow, where they regulate the permanent production and renewal of all blood-cell types. HSPC proliferation and differentiation is locally regulated by their interaction with cells forming specific microenvironments close to the bone matrix or close to blood vessels. However, the cellular mechanisms underlying HSPC’s interaction with these cells and their potential impact on HSPC polarity is still poorly understood. Here we modelled the bone-marrow niche using microfluidic technologies in a bone-marrow on a chip device, and evaluated long-duration cell-cell contacts between single HSPCs and stromal cells or endothelial cells in a custom-designed microwell cell-culture system. We found that an HSPC can form a discrete contact site that leads to the extensive polarization of their cytoskeleton architectures. As in the case with immune synapses formed by lymphocytes, the centrosome was located in proximity of the cell-cell contact. The entire microtubule network emanated from the centrosome, and the nucleus was confined to the side opposite of the cell-cell contact. The capacity of the HSPC to polarize appeared specific as it was not observed in contact with skin fibroblasts. The receptors ICAM, VCAM and CXCR4 were identified in the polarizing contact, and were all independently capable of inducing morphological polarization. However, only CXCR4 was independently capable of inducing the polarization of the centrosome-microtubule network. Altogether these results revealed a novel mechanism of HSPC polarization associated with its anchorage to specific cells in the bone-marrow, which might be instrumental in the regulation of their fate.

## Introduction

Hematopoietic stem cells are at the origin of all blood lineages (Orkin and Zon, 2008). In the liver of the fetus, or in the bone-marrow of adult, hematopoietic stem and progenitor cells (HSPCs) sense and respond to numerous biochemical stimuli (Pinho and Frenette, 2019). Within the bone-marrow, the vascular network and the bone matrix constitute local niches that impart distinct and specific signals regulating the quiescence, proliferation and differentiation of HSPCs (Morrison and Scadden, 2014) (Christodoulou et al., 2020) (Guezguez et al., 2013). Perturbed interactions between HSPCs and their niches have been associated with blood malignancies and ageing (Verovskaya et al., 2019), underscoring the importance of better understanding how HSPCs sense and respond to stromal and endothelial cells in the bone-marrow (Ceafalan et al., 2018).

Several lines of experimental evidence, in living organisms and in cultured cells have revealed that, in addition to diffusible signals, direct cell-to-cell contact is involved in the regulation of HSPC fate (Wagner et al., 2007)(Bruns et al., 2014)(Alakel et al., 2009)(Ceafalan et al., 2018)(Walenda et al., 2010). In co-cultures of human CD34+ HSPCs isolated from newborn cord blood and mesenchymal stromal cells from bone marrow aspirates (Wagner et al., 2007), HSPCs can adopt elongated and asymmetric morphologies, with several types of protrusions of various length and width that can have specific impact on proliferation and differentiation (Freund et al., 2006)(Frimberger et al., 2001)(Holloway et al., 1999). Similar polarized HSPC morphologies have also been observed in vivo (Coutu et al., 2017) but the stromal cells and signaling pathways that give rise to these morphologies remain to be deciphered.

In addition to a polarized morphology, HSPCs can polarize biochemically, as characterized by the accumulation of membrane-associated proteins in the protrusions forming either at the side in contact with the stromal cells (Freund et al., 2006) (Wagner et al., 2008) or at the opposite side (Görgens et al., 2012)(Fonseca et al., 2010). This polarization of membrane markers has been mostly described in the case of migrating HSPCs (Fonseca and Corbeil, 2011). Indeed, the segregated molecules and the associated signaling pathways are in many ways similar to the uropod of a migrating lymphocyte or neutrophil, and include the rearward localization of the centrosome (Sánchez-madrid and Serrador, 2009)(Fonseca et al., 2010; Heasman et al., 2010). However the uropod is also involved in cell-cell interactions in T lymphocytes, (Sánchez-madrid and Serrador, 2009) and HSPCs (Wagner et al., 2008), suggesting that not only migration but also anchorage could involve HSPCs polarization. In support of this hypothesis, localized adhesion-associated signaling and exchange of endosomes between HSPCs and osteoblasts have suggested the existence of synapse-like interactions, as it is the case for many stem cells interacting with the cells forming their niche (Wilson and Trumpp, 2006) (Ceafalan et al., 2018)(Gillette et al., 2009). However, the cellular mechanism inducing the polarization of HSPCs in response to their adhesion to stromal cells has not yet been investigated in detail. Furthermore, the similarities and differences between the polarities of migrating and anchored HSPCs are still unclear. Such investigation appear all the more necessary that it has recently been revealed that quiescent long-term hematopoietic stem cells are actually non-motile in vivo (Christodoulou et al., 2020). Although a lot has been learned from co-culture experiments, it has remained technically challenging to study the specific role of cell adhesion independently of cell migration.

## Results

In the bone marrow, hematopoietic progenitors encounter a large diversity of micro-environments, where diverse sets of stromal cells secrete specific cytokines and present specific inter-cellular adhesion receptors at their surface (Pinho and Frenette, 2019). In the endosteal niche, close to the bone matrix, osteoblasts interact directly with HSPC and thereby promote HSPC quiescence and long-term self-renewal capacities (Bowers et al., 2015)(Jung et al., 2005)(Guezguez et al., 2013)(Calvi et al., 2003). By contrast, in the peri-vascular niches, close to blood veins and arteries, endothelial cells and pericytes stimulate HSPC proliferation and differentiation (Kopp et al., 2005)(Kiel et al., 2005)(Ding et al., 2012)(Greenbaum et al., 2013)(Asada et al., 2017). To investigate the molecular and cellular mechanisms underlying these activities, we designed a microfluidic bone-marrow on a chip model of the HPSC niche (Ingavle et al., 2019)(Chou et al., 2020)(Sieber et al., 2018). The model was inspired by the pioneering work of Noo-Li Jeon, who described the set-up for the micro-channel geometry and the culture conditions necessary for inducing endothelial cells self-organization into hollow and perfusable 3D networks (Kim et al., 2013). Side channels included osteoblasts in 3D matrices made of collagen and fibrin to model a minimal version of the endosteal niche (Nelson et al., 2019). Maskless photo-lithography was used to test several chip prototypes and optimize channel design, which included the presence of pillars preventing the collapse of the 3D matrix in response to high contractile forces produced by osteoblasts (see Methods and Supplementary Figure S1). Human HSPCs (CD34+ from new born cord blood) were loaded in a central channel with the same 3D matrix as in the side channels (Figure 1A), so that they could migrate in 3D and enter those side channels (Figure 1B, Supplementary movie S1).

**Figure 1:**
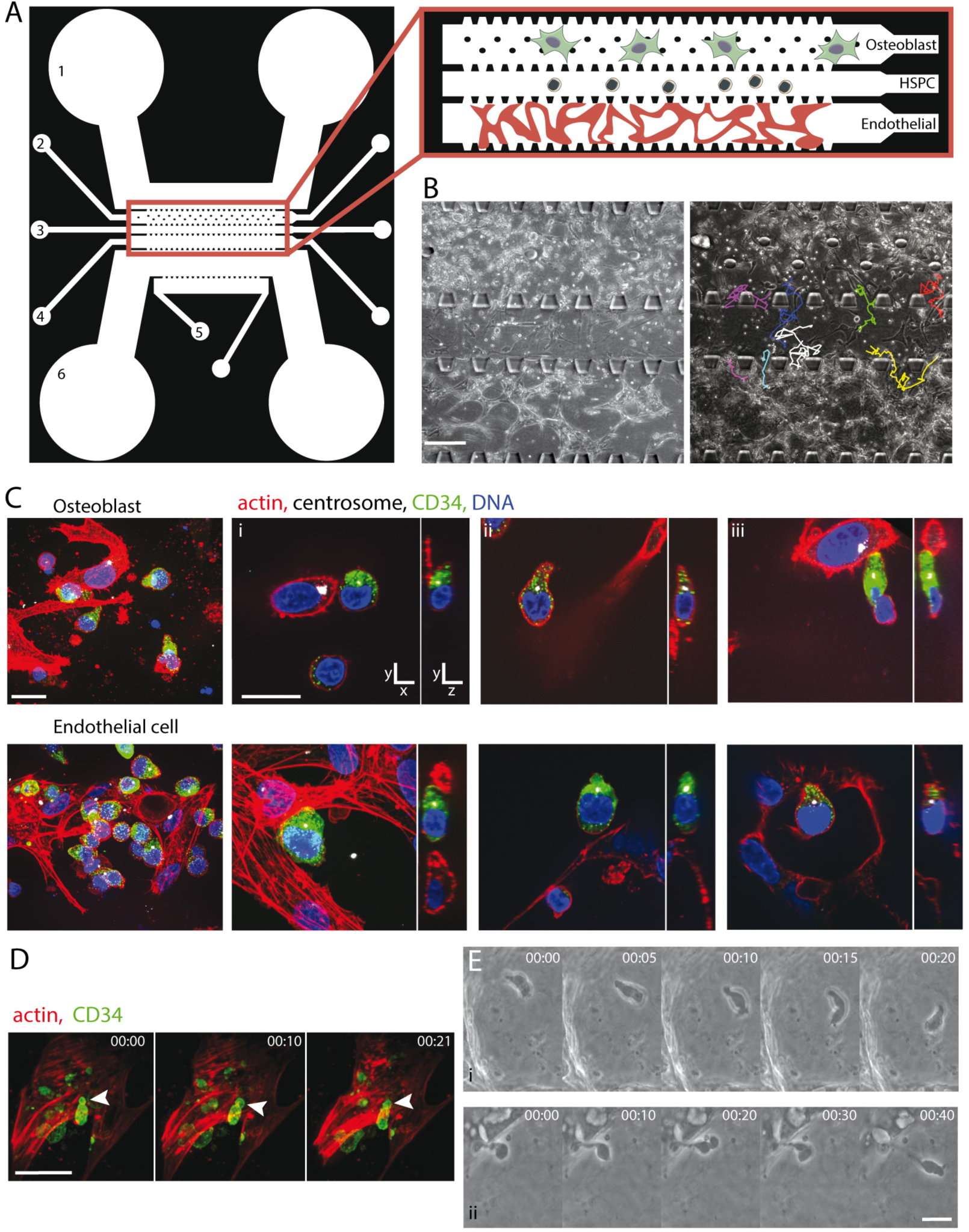
Bone-marrow on a chip allows the monitoring of HSPCs in contact with osteoblasts and endothelial cells in 3D hydrogels. (A) Illustration of the microfluidic chip design. The chip comprises channels for medium circulation (1 and 6), the endosteal compartment (2), the vascular compartment (4), the HSPC injection channel (3), and cytokine-secreting fibroblasts (5). The inset on the right describes the organization of the three central channels. (B) The three central channels of the chip in transmitted light (left image). The highlighted trajectories of several HSPCs during a time-lapse sequence (right image). Scale bar = 200 µm. (C) The shape and polarity of HSPCs revealed by confocal fluorescence microscopy in the chip. HSPC are revealed by CD34 staining (green). Actin filaments are shown in red, DNA in blue and the centrosome in white. Upper images show the endosteal compartment, and the lower images show the vascular compartment. Left images show the maximum projection of a 10 µm-wide z stack. The three composite images on the right show single xy-planes and respective single xz-planes illustrating elongated morphology of the HSPC and the centrosome positioning with respect to the contact site. Scale bar = 10 µm. (D) A time-lapse sequence (using confocal fluorescence microscopy) of an HSPC (CD34^+^) in contact with an osteoblast (Lifeact-stained actin filaments in red) revealing anchoring and deformation of an HSPC as it contacts the osteoblast. Scale bar = 20 µm. (E) A time-lapse sequence (transmitted light) of an HSPC in contact with osteoblast, showing (i) HSPC detachment and migration, and (ii) anchoring and deformation of the HSPC as it contacts osteoblast. Scale bar = 10µm.

Importantly, the bone-marrow on a chip model was compatible with chemical fixation, immuno-labelling and high magnification imaging, allowing actin networks and labelled centrosome to be visualized to reveal cell-shape polarization and the position of the main microtubule-organization center (MTOC). HSPCs were identified by CD34 staining. Cell-cell contacts were imaged in 3D to capture all orientations. In the bone-marrow on a chip model, HSPCs displayed both round and polarized shapes in contact with either osteoblasts or endothelial cells (Figure 1C), as found in vivo (Coutu et al., 2017),. All types of MTOC positioning were observed, either towards the site of contact or towards the opposite side (Figure 1C). However, their exact orientation with respect to the contact site was difficult to measure due to the frequent multiplicity of contact sites. Live imaging revealed that the contact between an HSPC and osteoblast or endothelial cells could be transient or last up to a few hours (Figure 1D). Furthermore, these contacts appeared strong enough to resist detachment due to cell migration in the fibrin hydrogel (Figure 1E). Although HSPCs displayed some clear and characteristic polarized organization, it was unclear whether this polarization was due to the cell-cell contact and not to the HPSCs high propensity to migrate in the 3D model.

To study the specific role of the contact between an HSPC and a bone-marrow niche cells, we developed a culture model that promoted long-term cell-cell interactions but prevented cell migration. Various sorts of microwells have been engineered to confine distinct cell types in a common volume (Dusseiller et al., 2005)(Khademhosseini et al., 2006)(Moeller et al., 2008)(Lutolf et al., 2009)(Guldevall et al., 2010)(Minc et al., 2011)(Gobaa et al., 2011)(Müller et al., 2015). In such microwells, HSPC stemness could be maintained over several weeks in 3D co-cultures with mesenchymal stromal cells (Wuchter et al., 2016). In our culture model, we used a differential patterning approach to restrict cell-substrate adhesion to the bottom of the microwell only in order to prevent cells from escaping the well (Dusseiller et al., 2005; Ochsner et al., 2007)(Gobaa et al., 2011) and to enable high quality imaging. This approach required a new fabrication protocol combining glass silanization and poly-acrylamide (PAA) capillary-based molding using non-adhesive microwells with glass bottoms (see Methods and Supplementary Figure S2).

HSPC were cultured at approximately one cell per 50-µm-wide microwell already seeded with a single osteoblast. As expected, HSPCs interacted with the dorsal surface of the osteoblast (Figure 2A). Long-term imaging showed that HSPCs proliferated at a normal rate, suggesting that the microwell manufacturing was not toxic (Figure 2B). Furthermore, HSPCs were occasionally observed to migrate, and locate below osteoblasts and proliferate, forming what has been termed a cobblestone structure that is typical of hematopoietic stem cells in long-term cultures (Jing et al., 2010) (Figure 2C), further suggesting that the HSPC were healthy. Certain HSPCs were observed to attach to migrating osteoblasts by video recording, showing that this co-culture model permitted the formation of strong heterotypic cell-cell contacts (Figure 2D) as previously found in other models (Wagner et al., 2007). Interestingly, HSPCs attached osteoblasts via a small but strong anchorage site that resisted cell migration despite dynamic shape changes (see supplementary movie S2). Attached HSPCs adopted elongated and asymmetric shapes (Figure 2E) similar to those observed in 3D conditions in the bone-marrow on a chip model (Figure 1E), as well as in bone-marrow in vivo (Coutu et al., 2017) and in other co-culture models (Freund et al., 2006). The contact site was restricted to a small area estimated to be around 1–2 µm^2^. However, HSPCs were not observed to spread on osteoblasts, in contrast to a lymphocyte forming immune synapse on a target cell (Ritter et al., 2013). Nevertheless, the cell architecture was highly polarized and similar to that of an immune synapse (Figure 3A). In particular, the centrosome was typically observed at the tip of the protrusion, in close proximity with the contact site (Figure 3A, and 3D reconstitution in supplementary movie S3). Considering that these HPSCs were not actively migrating, and sometimes orthogonal to the dorsal surface of the osteoblasts (Figure 3A), this polarized structure could not be confused with the uropod at the rear of a migrating lymphocyte. The architecture of polarized HSPC was characterized by accumulation of dense actin networks at the anchorage site and in a tail-like structure at the opposite side (Figure 3B). As stated above, the centrosome, the main if not the only microtubule-organizing center in these cells, was found in the protrusion associated with the contact site, and at a distance from the nucleus (Figure 3A, and 3C). Microtubules emanated from the centrosome, lined up along the cell membrane and all around the nucleus. In particular, microtubules accumulated in the wide cleft of the nucleus facing the protrusion, suggesting that microtubules were applying pushing forces responsible for the deformation of the nucleus (Figure 3C). No protrusion and no separation between the centrosome and the nucleus were observed when HSPCs were plated on a non-adherent surface (i.e. in a microwell coated with PAA; see Supplementary Figure S3).

**Figure 2:**
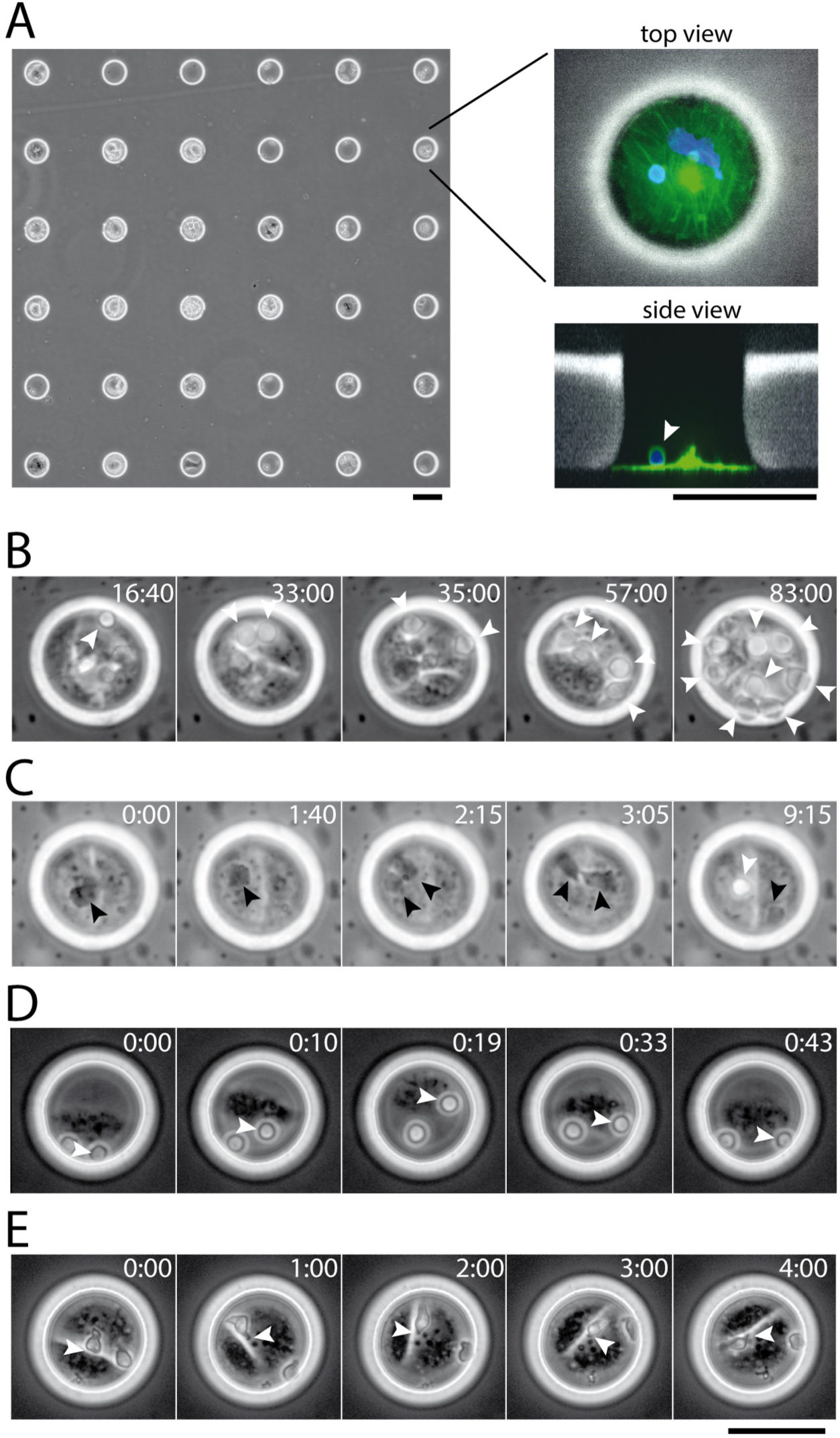
Array of microwells to control the interaction of HSPC with stromal cells. A) Images (in transmitted light) of the poly-acrylamide stencil showing the 50-µm wide circular holes seeded with osteoblasts and HSPCs (left). Images of top (upper right) and side (lower right) views of a single microwell containing fixed cells stained for tubulin (green) and DNA (blue). A fluorescent dextran is incorporated in the poly-acrylamide mix to reveal the microwell in fluorescence (white). An HSPC was identified by its small size and round shape (white arrowhead), whereas an osteoblast was larger and flatter in shape, and spread at the bottom of the microwell. (B-E) Time-lapse monitoring with transmitted light of live HSPCs (white arrow heads) in contact with osteoblasts (time indicated in hours:minutes), revealing; (B) the proliferation of an HSPC; (C) the migration and confinement of HSPC below osteoblasts (highlighted with black arrow heads), (D) the adhesion of HSPC onto a moving osteoblast; and (E) the long-term anchoring of an HSPC upon contact with an osteoblast, revealing the focused anchoring point and the asymmetric deformation of its shape (see corresponding Supplementary Movie S2). Scale bars = 50 µm.

**Figure 3:**
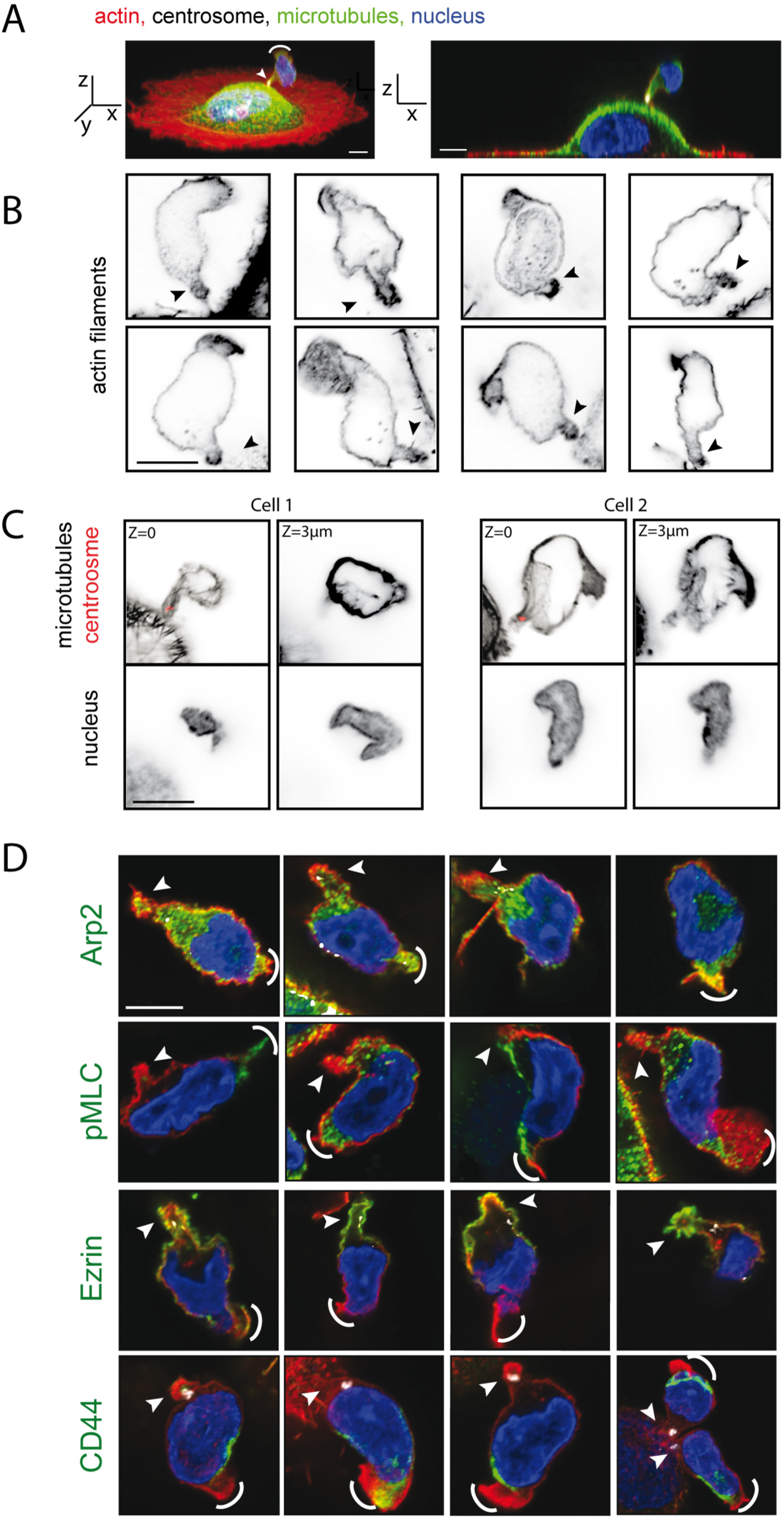
The polarized cytoskeletal architecture of HSPCs in contact with osteoblasts. (A) Representative confocal images (tilted 3D view, left; and lateral view, right) of an HSPC cultured with osteoblasts in a microwell for 15 h, fixed and stained for actin (red), microtubules (green) and centrosome (white). See supplementary movie S3 for a 3D reconstruction. The images show a polarized HSPC in which the centrosome is in close proximity to the site in contact with the underlying osteoblast. (B) Eight representative examples of polarized actin networks in fixed HSPCs in contact with osteoblasts. (C) Two representative examples of polarized microtubule networks in fixed HSPCs in contact with osteoblasts. Two z-stacks (z= 0µm and z=3 µm) are shown corresponding to the position of the centrosome (red, left) and the mid-section of the nucleus (right). (D) Four representative examples of fixed polarized HSPCs in contact with osteoblasts. Actin filaments are shown in red, centrosomes in white and DNA in blue. The four rows reveal Arp2, pMLC, ezrin, and CD44, respectively (green). A white arrowhead indicates the site of contact with the osteoblast and the white arc indicates the distal tail of HSPC. Scale bars = 5 µm.

HSPC polarization was further characterized using other classical markers of polarized compartments in lymphocytes, such as uropod or immune synapse (Sánchez-madrid and Serrador, 2009; Ritter et al., 2013). Arp2/3 and ezrin appeared concentrated in the protrusion situated at the contact site (Figure 3D). The phosphorylated form of myosin-II was observed in both protrusions situated at the contact site and at the other side of the cell (Figure 3E). CD44, a well-characterized uropod marker (Gomez-Mouton et al., 2001), was absent from the contact-associated protrusion but localized in the protrusion at the other side of the cell (Figure 3E). These markers suggested that the contact-associated protrusion differs from a uropod and more resembles an immune synapse despite some differences, such as its size and morphology. Altogether, these results showed that HSPC developed highly polarized cytoskeleton and membrane-associated architectures in response to forming a contact with an osteoblast. The similarity with the immune synapse raised the question of the specificity of the target cell and prompted us to test whether HSPC could polarize in contact with any type of stromal cell, and whether this polarization was an exclusive feature of a progenitor cell.

We thus examined the polarization of HSPCs (CD34+) on human umbilical vein endothelial cells (HUVEC), human bone-derived osteoblast (hFOB), human skin fibroblast (BJ) and murine liver-derived mesenchymal stromal cells (Figure 4A). As previously described, adherent cells were plated first and HSPCs were added after. Fifteen-hours later, cells were fixed and stained to assess cell shape and centrosome positioning. To quantify HSPC polarization towards the contact site, we measured the cell-polarity index, defined as the ratio between the centrosome distance to the contact site with respect to cell length (Figure 4B). A values close to 0 attested to a cell with the centrosome close to the contact site, whereas a value close to 1 attested to a cell with centrosome on the other side opposite of the contact site(Figure 4B). Interestingly, polarization was observed when an HSPC formed a contact with umbilical vein endothelial cell and osteoblast, but not when in contact with a skin fibroblast (Figure 4C). To assess whether polarization was related to a functional role that stromal cells have on HSPC regeneration potential, we compared the polarization of HSPC with two murine fetal-derived stromal cell lines; AFT024, which is known to support HSPC regeneration capacities ex vivo, and BFC012, which is not (Moore et al., 1997)(Charbord et al., 2014). In support of this notion, HSPCs polarized only when in contact with AFT024 cells (Figure 4D). We further assessed the selectivity of the polarizing interaction, in terms of the differentiation status of the hematopoietic cell by comparing hematopoietic stem cells (HSC; CD34+/CD38 low), common myeloid progenitors (CMP; CD34+/CD38 high/CD33 high; see Supplementary Figure S4 for parameters of FACS sorting) and mature T lymphocytes (Jurkat cells) in contact with osteoblasts. We found that for both types of progenitor cells (HSCs and CMPs) when in contact with osteoblasts, marked polarization was observed (Figure 4E). By contrast, for Jurkat cells, no polarization was observed (Figure 4E), even though Jurkats display distinct centrosome polarization when in contact with antigen presenting cells (Yi et al., 2013). All together, these results showed that the hematopoietic polarization is not a generic outcome but specific to defined interactions between hematopoietic progenitors and stromal cells. The results also suggested that when in contact with stromal cell, a stem cells is more likely to polarize than a differentiated cell, and that the capacity to induce polarization of an HSPC is greater for a stromal cell from the bone-marrow niche than other stromal cells. Hence, the polarization of an HSPC by the stromal cell may be induced by defined combination of surface ligands and receptors.

**Figure 4:**
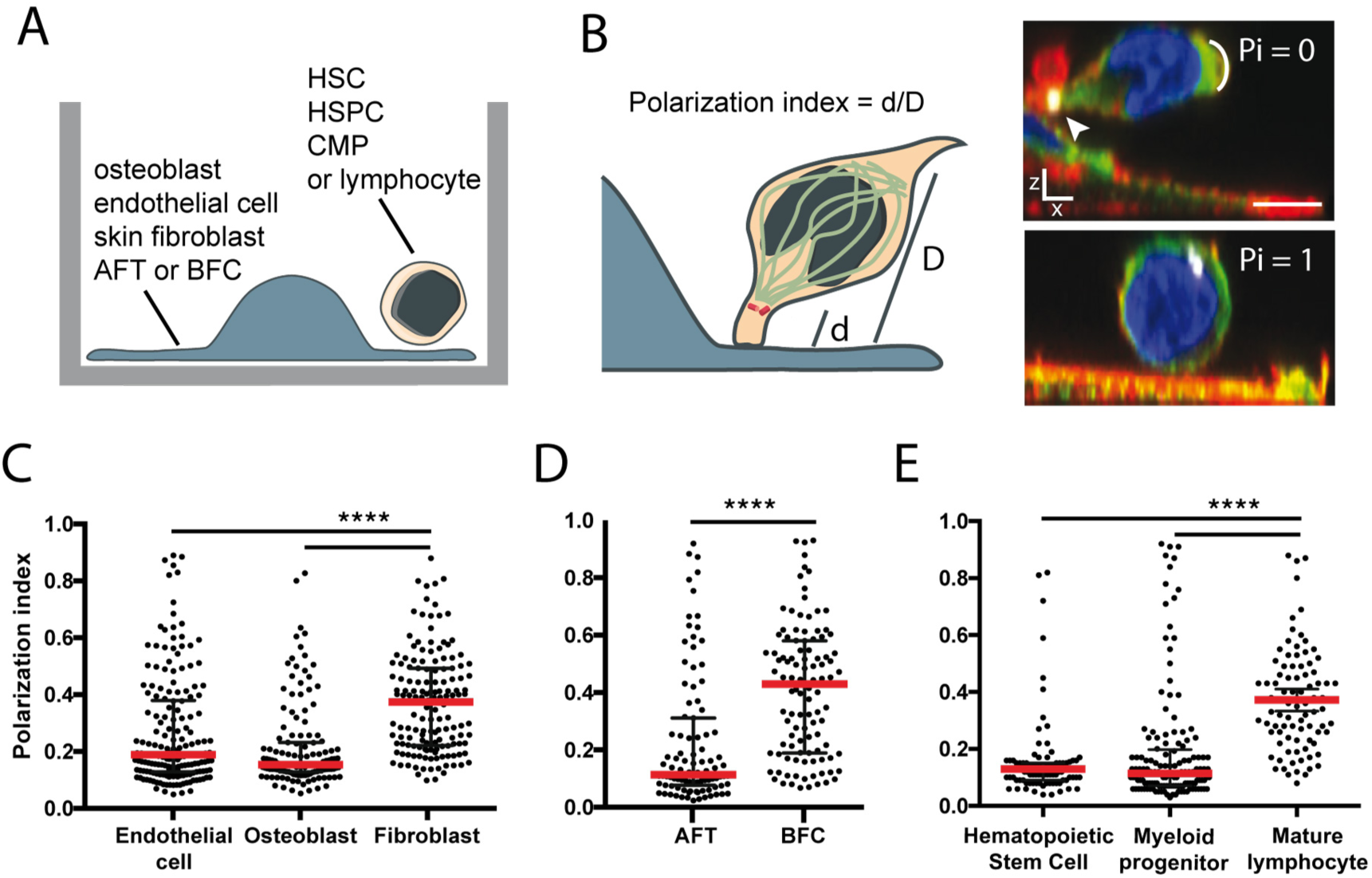
Polarization depends on specific heterotypic interactions between the HSPC and stromal cells. (A) Schematic description of the experimental strategy to evaluate interactions between different cell types. Different hematopoietic cells populations, from stem to fully differentiated cells, were seeded on different stromal cell types for 15 hours and fixed. (B) HSPC polarization was defined by the distance, d, between the position of the centrosome and the point of contact with the stromal cell, divided by the cell length, D, from that point of contact. Representative images of HSPCs with either a polarization index (Pi) close to 0 (upper right), or close to 1 (lower right). Actin filaments are in red, microtubules in green, the centrosome in white and DNA in blue. Arrowheads highlight the protrusion in contact with the stromal cell and the arc indicates the distal tail of the HSPC. Scale bars = 5 µm. (C to E) Scatter plots of polarization indices of (C) HSPCs in contact with endothelial cells, osteoblast or skin fibroblast; (D) HSPCs (CD34^+^) in contact with liver-mouse-derived stromal cell lines that either support HSPC regeneration capacities (AFT024) or not (BFC012); and (E) hematopoietic stem cells (CD38^-^/CD34^+^), common myeloid progenitors (CD34^+^/CD33^+^) or mature lymphocytes in contact with osteoblasts. Median and interquartile range indicated by red and black horizontal bars, respectively. **** = p value < 0.0001.

Several pathways are involved in the physical interaction and biochemical crosstalk between hematopoietic progenitors and niche cells (Wilson and Trumpp, 2006)(Ceafalan et al., 2018). To identify those that were involved in the polarization of HSPC, we immunolabelled receptors known to play key roles in cell adhesion and the regulation of hematopoietic differentiation (Ceafalan et al., 2018), including the receptor pairings, VCAM-VLA4, ICAM-LFA1, and SDF1-CXCR4. All receptors appeared to be polarized and localized in the protrusion associated with the contact site of HSPC with the osteoblast (Figure 5A). To investigate the independent impact of a specific pathway, HSPCs were seeded into microwells, the bottoms of which were coated only with ligands of a particular receptor, (Figure 5B). Live monitoring showed that HSPCs attached to the bottom of the microwell and adopted the same type of elongated and polarized shapes that were observed for the contacts with osteoblasts (Figure 5C). This was observed with the ligands, SDF-1, ICAM-1 and VCAM-1, but not with the negative controls of non-adherent poly-acrylamide or no coating (Figure 5D). Strikingly, the centrosome and microtubules were polarized only in the HSPC in contact with SDF-1 (Figure 5E), and not with ICAM-1 or VCAM-1, or with poly-acrylamide or no coating (Figure 5E). These results show that although all receptors appeared engaged in the polarization process (by localization) only SDF-1 appeared sufficient to autonomously induce the morphological and internal polarization of the HSPC.

**Figure 5:**
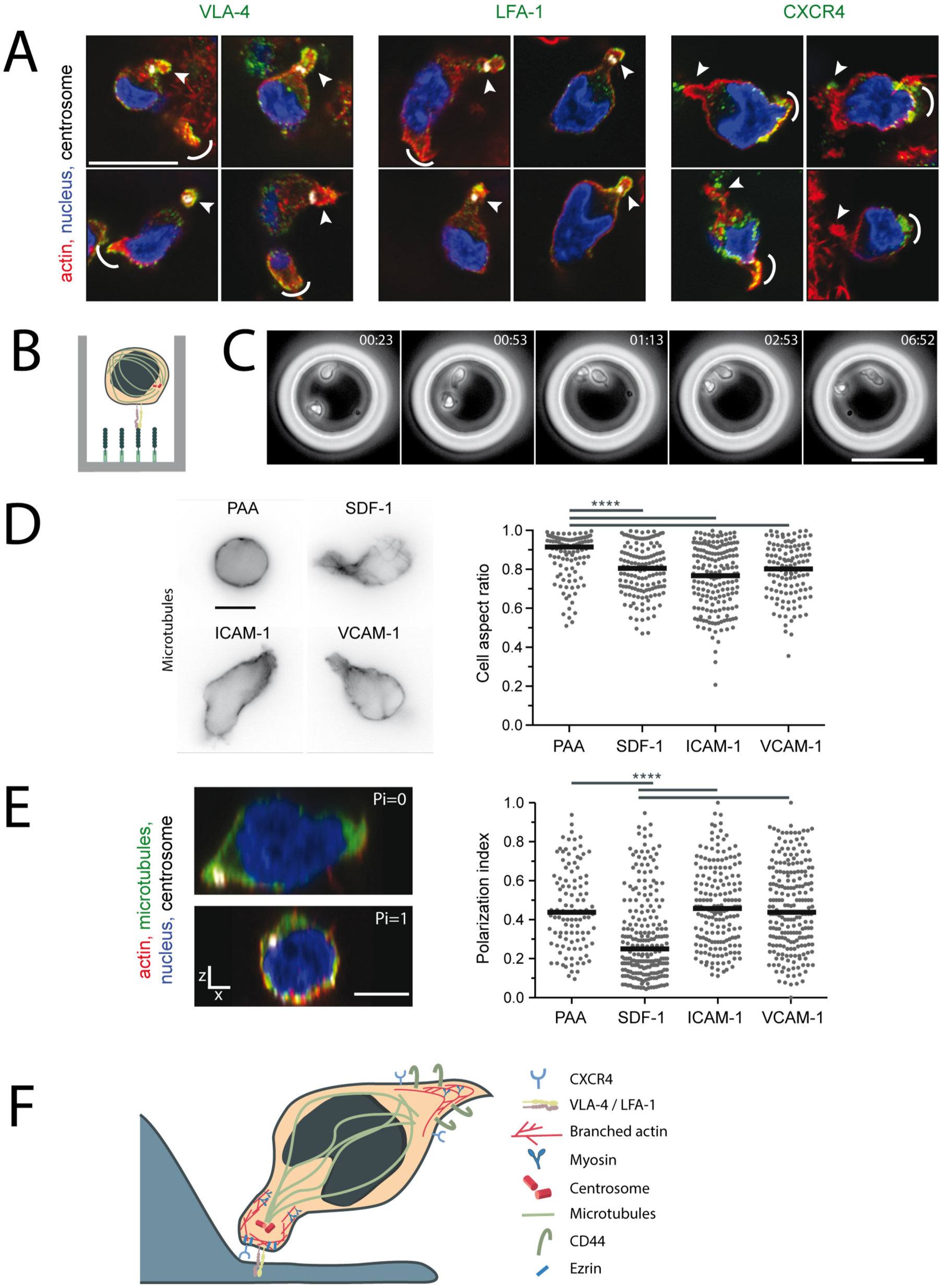
Engagement of CXCL12/CXCR4 is sufficient to induce HSPC polarisation. (A) Representative confocal images of an HSPC cultured in a microwell for 15 h, fixed and stained for actin (red), microtubules (green) and centrosome (white). HSPCs cultured with osteoblasts in microwells for 15 h, fixed and stained for actin (red), DNA (blue) and centrosome (white). Three panels of four examples of cells also stained in green for CD49d/VLA-4 (left), CD18/LFA-1 (middle) or CXCR4 (right). Arrowheads highlight the protrusion in contact with the osteoblast and the arc indicates the distal tail of the HSPC. Scale bar = 10 µm. (B) Schematic illustration of an HSPC in a microwell coated with protein A (light green) and Fc-tagged-protein with the potential to function as a ligand for an HSPC receptor (dark green). (C) Time-lapse sequence (transmitted light) of HSPCs in a microwell coated with SDF-1. Time annotation is in h :min. Scale bar = 50 µm. (D) Representative images of HSPCs (left panel) cultured on SDF-1, ICAM-1 or VCAM-1 coated microwells or uncoated microwells (PAA), respectively. The morphology of the HSPC is revealed by microtubule staining. (Right panel) Scatter plot of the cell-aspect ratios of the HSPCs cultured in the different coated microwells. The cell aspect ratio was calculated as the ratio of the lengths of the short and long axes of the cell. (E) Representative images of an HSPC cultured on an SDF-1–coated microwell with a polarization index (Pi) close to 0 (upper left), and an HSPC cultured on an uncoated microwell with a Pi close to 1 (lower left). Actin filaments are in red, microtubules in green, the centrosome in white and DNA in blue. (Right panel) Scatter plot of polarization indices of HSPCs cultured in the different coated microwells, as in (D). Median and interquartile range indicated by red and black horizontal bars, respectively. **** = p value < 0.0001. Median and interquartile range indicated by red and black horizontal bars, respectively. **** = p value < 0.0001. (F) Schematic representation of the shape and polarized architecture of an HSPC in contact with a stromal cell.

## Discussion

In modelling the bone-marrow niche in vitro, we have identified novel cytoskeletal architectures and molecular signatures characterizing the interaction and polarization of the HSPC when it forms a contact with stromal or endothelial cells. Our results differ from what has been described previously for the morphological polarization of an HSPC undergoing migration, which, at its rear edge, assembles a protrusion that shares many features with the uropod of a migrating lymphocyte or neutrophils (Fonseca et al., 2010)(Görgens et al., 2012). With the formation of a long-lasting contact with osteoblast, the hyaluronic acid receptor CD44, which is a characteristic marker of the lymphocyte and HSPC uropod (Gomez-Mouton et al., 2001)(Wagner et al., 2008), was not localized at the anchorage site, but in the protrusion at the other side of the cell (Figure 5F). In addition, the pointed morphologies of the protrusion and the small size of the anchorage point were other marked differences (Figure 5F). Interestingly, although ICAM, VCAM and CXCR4 were all segregated in the protrusion at the contact site with the osteoblast, ICAM- and VCAM-mediated adhesions appeared only to have the capacity to induce a morphological polarization of HSPC, whereas CXCR4 engagement with its ligand SDF-1 appeared also to have the capacity to induce the recruitment of the centrosome at the contact site. However, considering that CXCR4, ICAM, VCAM and other adhesion receptors mutually activate each other (Peled et al., 2000)(Glodek et al., 2007) (Petty et al., 2009)(Chang et al., 2016) it is likely that the complete molecular mechanism inducing and establishing the entire internal polarization of HSPC involves the synergy of several signaling pathways associated with the adhesion of the HSPC to the bone-marrow niche cells.

SDF-1 (CXCL12), the ligand that binds to CXCR4, is a major regulator of several key features of HSPC function, including the chemotactic mobilization towards the vascular niche and the maintenance of the pool of HSPCs (Lévesque et al., 2003)(Crane et al., 2017)(Greenbaum et al., 2013). It is interesting to consider that the polarization and anchoring we identified here (Figure 5F) could be involved in the homing and tethering of HSPCs to a particular aspect of the bone-marrow niche. The centrosome polarization close to the contact site is reminiscent of the structure of immune (Stinchcombe et al., 2006)(Ritter et al., 2013) and of the polarization of several other types of stem cells with their niches (Ceafalan et al., 2018). Whether such polarization of the HSPC by CXCR4 leads to local exchange of signaling molecules (Gillette et al., 2009) and structural reorganization that regulates the quiescence and/or the asymmetry of subsequent HSPC divisions (Ho and Wagner, 2007) are interesting possibilities that deserve further investigations.

## Acknowledgements

We thank Noo-Li Jeon and Dorian Obino for providing chips and tips for the vasculogenesis on chip model.

## Authors contributions

T. Bessy performed most experiments with the help of L. Faivre, B. Vianay, A. Schaeffer and S. Brunet. B. Souquet performed experiments in the bone-marrow on a chip model with the help of B. Vianay and S. Brunet. T. Jaffredo provided fetal liver cell lines and advice on the project. L. Blanchoin, J. Larghero, M. Théry and S. Brunet supervised the project. J. Larghero and M. Théry obtained funding for the project. M. Théry and S. Brunet conceived and directed the project. T. Bessy and M. Théry wrote the manuscript which was further critically reviewed by all authors.

## Materials and Methods

### Cells and culture

Human umbilical cord blood samples were obtained from the Cord Blood Bank of the Saint-Louis Hospital (France), in accordance with French national law (Bioethics Law n° 2011-814) and under declaration to the French Ministry of Research and Higher Studies. Using lymphocyte-separation medium (Eurobio), mononuclear cells were separated from erythrocytes and plasma by density gradient. CD34^+^ HSPCs were separated from other cells by magnetic sorting (MACS), using CD34 antibodies coupled with magnetic beads (Miltenyi Biotech). Cells were used directly after isolation. The cell lines used were; the human femural osteoblast line, hFOB (ATCC - CRL-11372), cultured in DMEM-F12 (Gibco); the human skin fibroblast line, BJ (ATCC - CRL-2522), cultured in αMEM (Gibco) and the normal human lung fibroblast line, NHLF (Lonza - CC-2512), cultured in FGM-2 (Lonza). AFT024 and BFC024 immortalized mesenchymal stromal cell lines were obtained from Lemishka’s group (Moore et al., 1997), and were cultured on gelatin-coated culture plates in DMEM (Gibco) with 50 µM β-mercaptoethanol. The human umbilical vein endothelial cell line, HUVEC (Lonza - 191027), was cultured on gelatin-coated culture plates in EGM-2 (Lonza). All medium were supplemented with 10% FBS and antibiotics/antimicotic (Sigma), except EGM2, that was supplemented with the EGM2 bulletlkit (Lonza).

### Flow Cytometry

Cells were stained for 30 min at 4°C in 500 µL of phosphate buffer saline (PBS) with 2 mM EDTA. Antibodies CD45-AF700 (BioLegend), CD38-PerCp5.5 (BioLegend), CD34-APC (BD Bioscience), CD33-PE (BD Bioscience), CD19-FITC (BD Bioscience) were used at 5 µL/10^6^ cells. The sorting procedure was performed on a FACS Aria II with DIVA software (BD Bioscience). After sorting, cells were centrifuged and resuspended in the desired volume of IMDM to achieve the appropriate cell-culture density.

### SU8 Mold Fabrication

Microwell shape, size and arrangement were drawn using the software CleWin, and a transferred onto a quartz photomask (Toppan) A wafer with micro-structures was made on glass (for the microfluidic chips) or on silicium (for microwells). Wafers were coated with a 5 µm layer of resin (MichroChem - CTS - SU8-3005). This layer was then fully exposed with UV light at 23 mJ/cm^2^ (Kloé - UV KUB2) for 5 s for full polymerization. Another layer of resin of 50 µm (MichroChem - CTS - SU8-3050) was spincoated on top of the first layer, for microfluidic chips this step was repeated a second time. This layer was exposed under the quartz mask with 23 mJ/cm^2^ UV light for 8 s for microwells, or with the PRIMO (Alveole) 32 mJ/cm^2^ for microfluidic chips. After development (MichroChem - CTS - Developer SU8) only the exposed structures remained. They were then hard baked for 2 h at 150°C, and coated with gas-phase trichloro(perfluorooctyl)silane (Sigma). For microfluidic chips, the glass wafer was countermolded with a silicone elastomer, base 9:1 crosslinker (Dow Corning – Sylgard 184 kit, PDMS), and is thereafter referred to as PDMS chip. For microwells, a negative mold of the silicium wafer was made with PDMS, it was then silanized in the same manner as the wafer. A second, positive mold of PDMS was made of the first mold, it is thereafter referred as PDMS stamp.

### Microfluidic

PDMS chips were punched in the circular openings, and then plasma bounded to glass coverslips. A CollFib hydrogel was made of thrombin (Sigma - T6884) 1 U/mL, fribrinogen (Sigma - F3879) 2 mg/mL, rat tail collagen-I (Ibidi - 50201) 1.6 mg/mL. 2Í10^5^ HUVEC and 1.5Í10^5^ hFOB were separately suspended in 20 µL of CollFib hydrogel, and loaded in their respective channels in the chips immediately after thrombin addition. NHLFs, previously treated with mitomicyn C, were suspended in 20 µL of thrombin 1 U/mL and fibrinogen 3 mg/mL, and injected in the chip. The chip was incubated for 30 min at 37°C. Osteoblast and endothelial medium were loaded in the large channels adjacent to their respective cell type. After 72 h of culture, CD34^+^ HSPCs in CollFib hydrogel were loaded in the central channel. The system was fixated after 4 d of coculture.

### Microwells

Thoroughly washed glass coverslips were plasmatized for 3 min, coated with gas-phase 3-(trimethoxysilyl)propyl methacrylate (Sigma), and baked at 120°C for 1 h. Coverslips were washed with ethanol before use. A PDMS stamp was plasmatized for 30 s and immediately placed on a silanized coverslip. Freshly-made solution of 20% acrylamide 37.5/1 bisacrylamide (Euromedex) in MiliQ water, with 1% APS and TEMED (Sigma) and 1% of photoinitiator (2-hydroxy-2-methylpropiophenone - Sigma) was immediately introduced by capillary action between the PDMS stamp and the glass coverslip. The sample was exposed to 23 mJ/cm^2^ UV light for 5 min. After exposure, the PDMS stamp was removed in MiliQ water.

Before use, microwell coverslips were coated with 40 µg/mL of protein in a 8.4 mg/L NaHCO_3_ solution for 15 min. Coverslips were coated with fibronectin (Sigma) for the plating of feeder cells, or sequentially with protein A (Interchim) and tag Fc proteins, SDF1-Fc, ICAM-1-Fc, or VCAM-1-Fc (Interchim), for functionalized wells for exposure to HSPCs alone. Chips were kept overnight in PBS before use for salt-equilibrium and photoinitiator detoxification. Chips were rinsed twice in medium immediately before use. For cell-cell interactions, 15000 feeder cells were seeded into each well; and to ensure that the cells entered into the wells, the chip was centrifuged. The chips were incubated for 1 h to promote cell spreading. Then, 15000 HSPCs were seeded over the wells and the chip was centrifuged again. For cell-protein interactions, 15000 HSPCs were seeded into each well, and the chip centrifuged.

### Immunofluorescent Staining

Cells were fixed for 15 min, after 4 d of culture for microfluidic chips, and after 15 h of culture for microwells, in cytoskeleton buffer (10 mM MES pH 6.1, 138 mM KCl, 3 mM MgCl, 2 mM EGTA, 10% sucrose) in 2% paraformaldehyde (PFA - Sigma) and 0.1% glutaraldehyde (Sigma) for cytoskeleton staining, or in 4% PFA for other antibodies. For phosphorylated myosin light chain (pMLC), cells were permeabilized 30 s in 0.5% triton in cytoskeleton buffer. Cells were permeabilized 10 min in 0,1% triton, except for surface markers where permeabilization was performed after the primary antibody. Coverslips were neutralized with a solution of NaBH_4_ (Sigma) for 10 min. The following primary antibodies and respective dilutions were used; rat anti-YL1/2 (ABD serotech), 1:500; rabbit anti-pericentrin (Abcam), 1:1000; anti-CXCR4 or anti-CD184 (BD Biosciences), 1:500; anti-CD18 clone TS1/18 (Thermo Fisher Scientific), 1:500; anti-CD49d (integrin α4) clone 9F10 (Thermo Fisher Scientific), 1:500; anti-Arp2 (Abcam), 1:500; anti-pEzrin, 1:200; anti-pMLC, 1:50; and anti-CD44 (R&D Systems), 1:500. Cells were incubated with secondary antibodies at 1:500, and where appropriate, phalloidin (Sigma). Finally, cells were incubated with DAPI (Sigma) for 10 min, and the coverslips were mounted with Mowiol (Sigma).

### Microscopy

Images for the quantification of centrosome polarization in HSPCs were captured on a upright Olympus BX61, wide field illumination, and with a CoolSnap HQ2 camera (Photometrics), and realized with Metamorph software. Immunofluorescence images were captured with a Nikon Ti-eclipse equipped with a spinning disk (Yokogawa - CSU-X1) and an EMCCD camera (Photometrics - Evolve 512) or a Retiga R3 (Q Imaging), and realized with Metamorph software. Illustration images were captured with a Zeiss LSM 800 microscope equipped with the Airyscan technology, and realized with ZEN software. For time-lapse imaging, cells were placed under the microscope immediately after seeding. Images were captured using a IX-83 Olympus microscope with an Orca flash 4.0 Lite camera (Hamamatsu) and realized with MicroManager software.

### Quantification

In the 3D images of the HSPC, three positions were identified; (i) the position of the centrosome (done automatically). (ii) the point on the cell membrane, which was nearest to the centrosome and was at the zone of interaction with the feeder cell (or the glass for HSPC-centrosome polarization on functionalized wells; done manually); and (iii), the point on the cell membrane, which was furthest from point (ii), excluding thin membrane protrusions (i.e. only including the main body of the cell). Distance d was defined as the length between points (i) and (ii) and distance D’ the length between points (ii) and (iii). The polarisation index was calculated as d/D.

All data are shown as scatter plots with median and interquartile range represented. A significance difference between populations was assessed with a non-parametric (Kruskal-Wallis) ANOVA test.

---

**Supplemental Figure S1:**
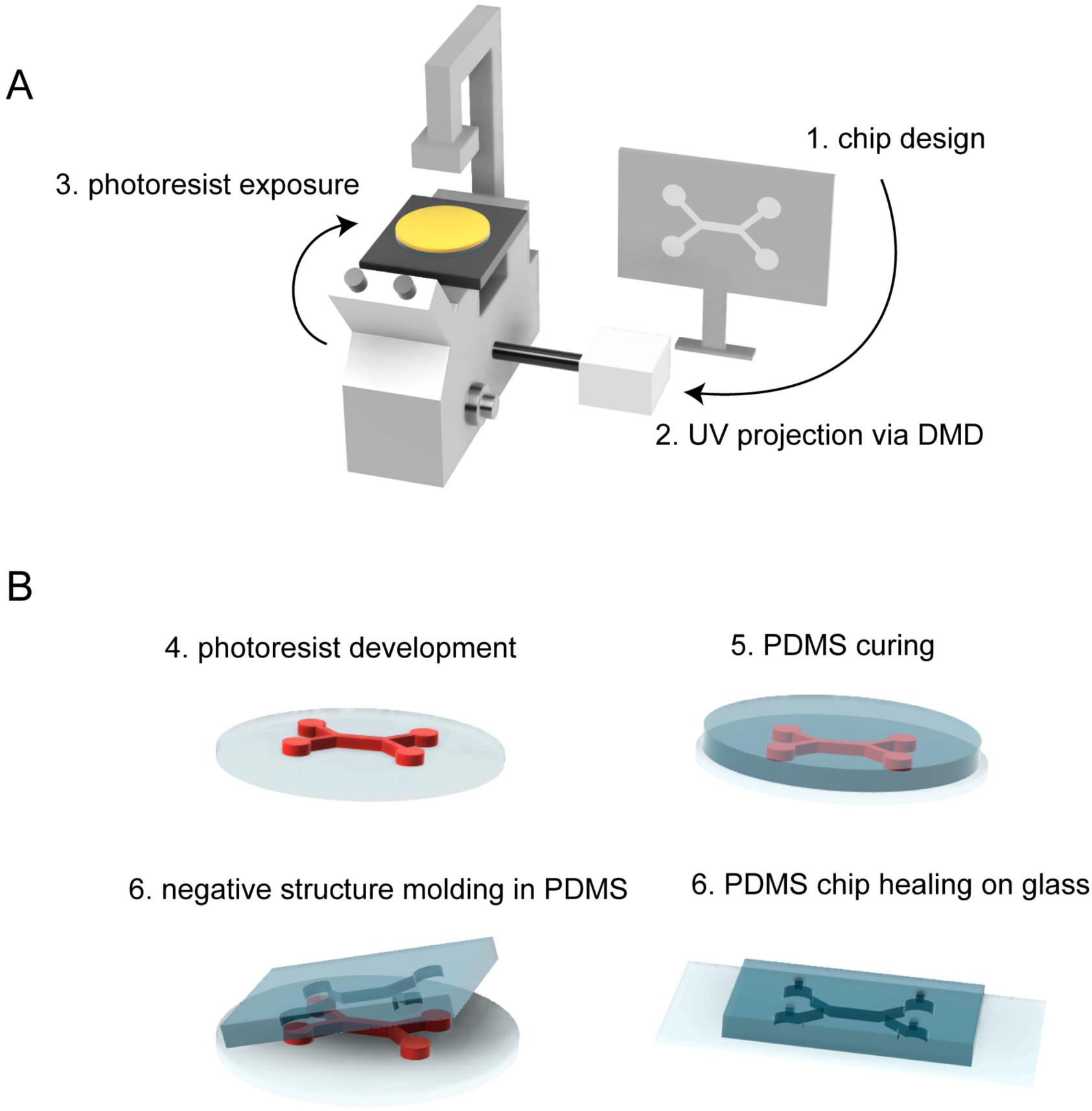
Microfluidic chip prototyping. (A) A photoresist was spun onto a glass wafer and exposed to UV via a digital micro-mirror device (DMD) with a chip geometry previously designed on computer. (B) The exposed photoresist was incubated in the developer, wash and bake. PDMS was further poured and cured onto the microstructures. The PDMS layer was then removed, put in contact with a glass slide and punched to plug flow inlets. The entire process, including the coating with the photoresist-layer and precuring, was performed in less than 1 h.

**Supplemental Figure S2:**
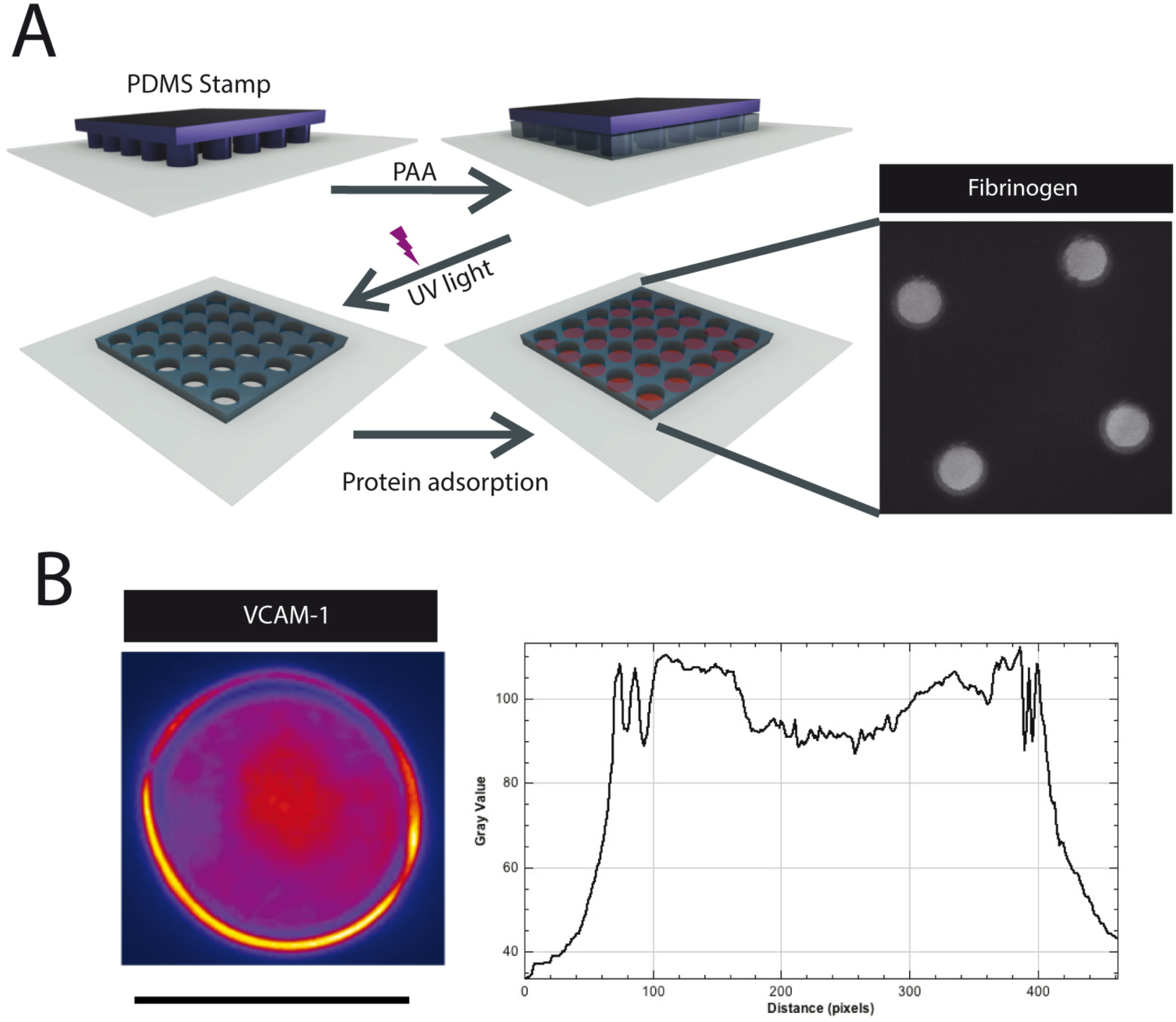
Microwell manufacturing. A) A polydimethylsiloxane (PDMS) stamp, manufactured with the same process as the one described in Figure S1 for microfluidic chips, was placed in contact with a silanized glass coverslip. A mixture of acrylamide and bis-acrylamide was introduced into the gap between the PDMS and the glass coverslip by capillary action. The sandwich was exposed to UV for 5 min to polymerize the poly-acrylamide (PAA). The PDMS mold was removed to reveal the stencil comprising open PAA microwells with glass bottoms. The glass bottom was functionalized by binding proteins to the silane as illustrated for fluorescent fibrinogen in the image on the right. (B) A representative image of VCAM-1 staining at the bottom of a PAA microwell. The linescan graph shows the fluorescence intensity across the microwell. Scale bar = 50 μm.

**Supplemental Figure S3:**
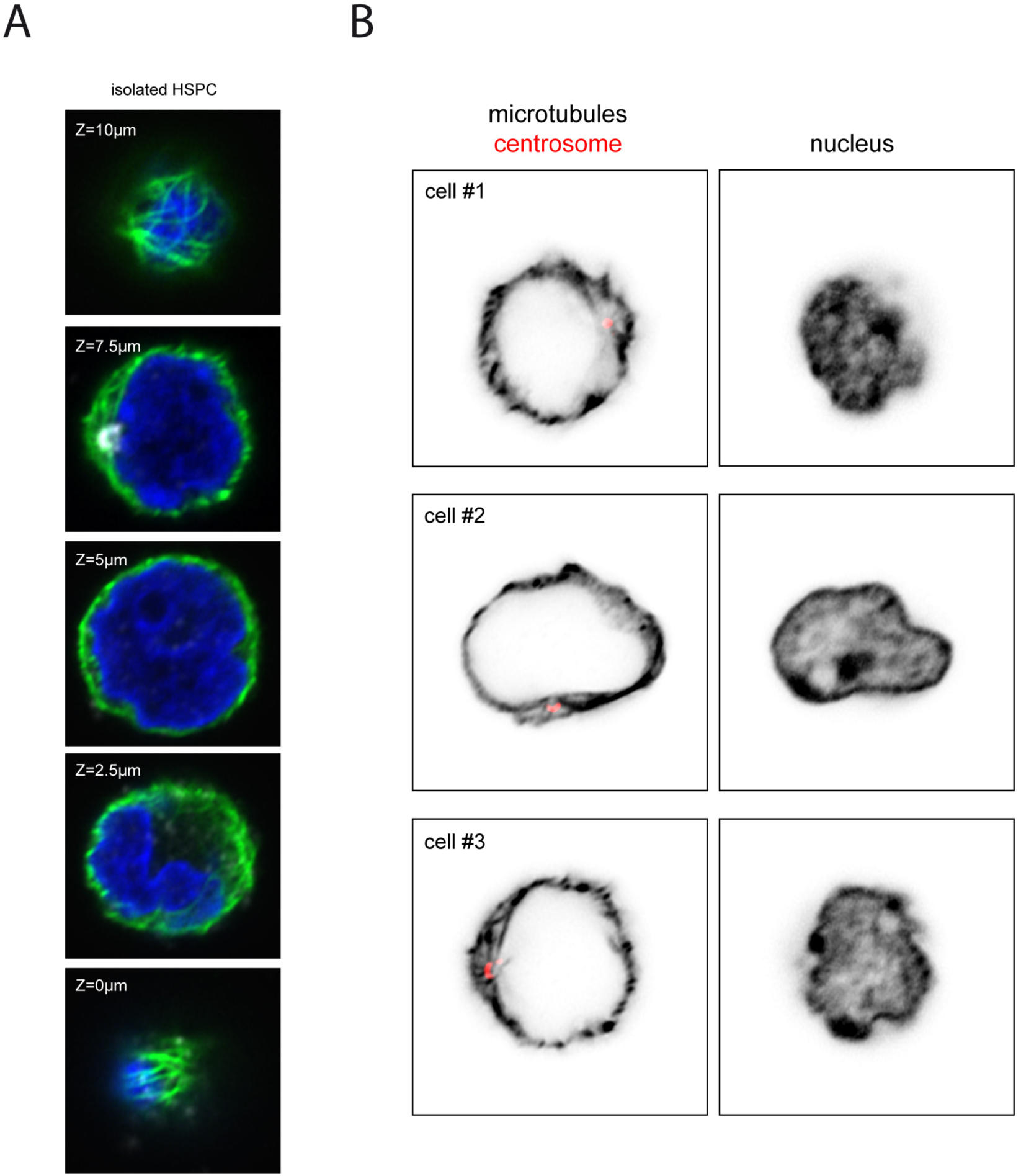
Cytoskeleton architecture of HSPCs in a non-adhesive microwell. Non-adhesive microwells were manufactured by pressing a PDMS stamp against a layer of acrylamide mix prior to polymerization. After UV exposure, this led to the formation of PAA microwells. (A) Representative confocal z-stack images (z=0–10 µm) of an HSPC cultured in a PAA microwell, and stained for microtubules (green), DNA (blue) and centrosome (white). The centrosome was not in contact with the glass bottom and close to the nucleus. (B) Three representative HSPCs cultured in PAA microwells stained for (left images) microtubules (gray) and the centrosome (red), and stained for (right images) DNA.

**Supplemental Figure S4:**
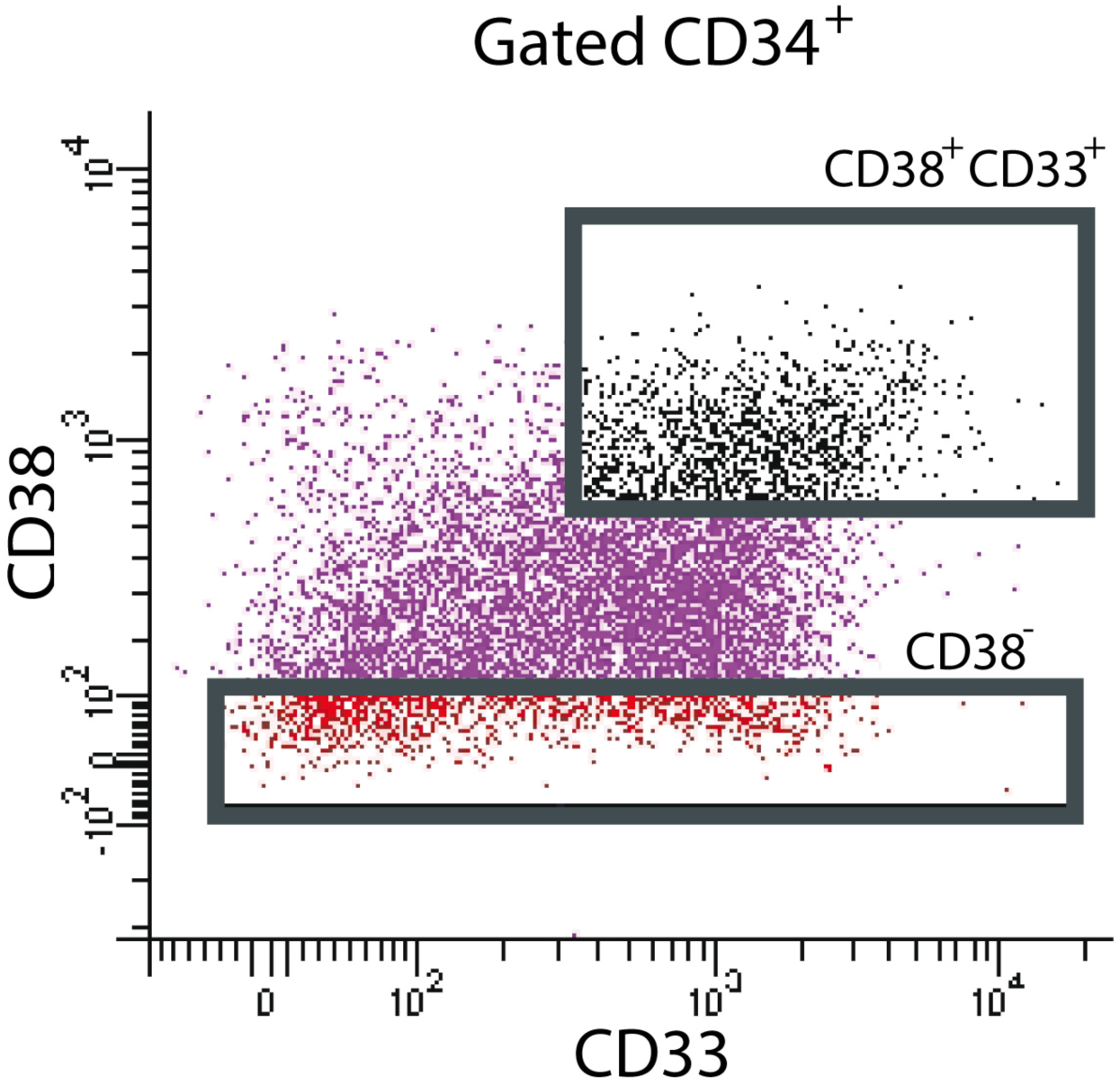
FACS sorting of HSC, HSPC, and CMP. Flow-cytometry output showing the magnetically-sorted separation of HSPCs. CD34^+^ cells were immunostained with fluorescent primary antibodies against CD34, CD38 and CD33. Cells were then further sorted to get CD34^+^/CD38^-^ cells, considered as HSC, and CD34^+^/CD38^+^/CD33^+^ considered as common myeloid progenitors.

## Supplemental movies

### Movie S1: HSPC migration in a bone-marrow-on-a-chip

Movie shows a transmitted light (phase contrast) video-recording of HSPC loaded in the central channel and migrating toward the pseudo-endosteal compartment (top channel) containing osteoblasts and the pseudo-vascular compartment (bottom channel) containing endothelial cells forming a vascular network. The pitch of central pilar spacing is 200 µm. Time is indicated in hours.

### Movie S2: HSPC anchorage to osteoblasts in a microwell

Movie shows a transmitted light (phase contrast) video-recording of two HSPCs on top of two osteoblasts in a microwell (50 micron-wide). Note the dynamic shape changes of HSPCs but their long-lasting anchorage on the dorsal surface of osteoblasts via a thin protrusion. Scale bar represents 20µm. Time is indicated in hours:minutes.

### Movie S3: HSPC polarization in contact with an osteoblast

Movie shows the rotation of a 3D reconstruction of a Z stack. It shows a single HSPC on top of an osteoblast in a microwell. Microtubules are shown in green, actin filaments in red and centrosomes in white. In the HSPC, note the position of the centrosome at the tip of the protrusion forming the contact with the osteoblast. The microwell diameter, and thus the width of the osteoblast, is 50 µm.

